# Multiplexed pooled library screening with Cpf1

**DOI:** 10.1101/473959

**Authors:** Jintan Liu, Sanjana Srinivasan, Chieh-Yuan Li, I-Lin Ho, Gang Wang, Wantong Yao, Angela Kay Deem, Chris Bristow, Traver Hart, Giulio Draetta

## Abstract

RNA interference and CRISPR/Cas9-based pooled library screens have revolutionized the field of functional genomics. However, currently available pooled library screens face a trade-off between library effectiveness and library complexity. We developed a multiplexed, high-throughput screening strategy based on an optimized AsCpf1 nuclease that minimizes library size without sacrificing gene targeting efficiency. Our AsCpf1-based multiplexed library performed similarly well compared to currently available CRISPR/Cas9 libraries, but with a single polycistronic crRNA clone targeting each gene. With this strategy, we constructed the smallest whole-genome knock-out library available, “Mini-human” for the human genome, which is one-fourth the size of the smallest CRISPR library currently available.

## Main

High-throughput forward genetic screens are an invaluable tool to systematically explore genetic interactions and to link gene disruption with disease contexts ^1^. The adaptation of CRISPR/Cas9 has resulted in pooled-library knockout screens with improved sensitivity and specificity versus pooled-library shRNA screens, primarily owing to more limited off-target effects, more potent gene perturbation, and higher proportion of active constructs in such system compared to an equivalent shRNA pool ^2–4^.

Cas9-based pooled library screens have demonstrated their feasibility and efficacy in numerous studies ^1,3,5^. Like pooled shRNA screens, pooled sgRNA screens benefit from multiple constructs targeting each gene of interest, as the functionality of each construct is unknown. However, increasing the number of genetic reagents per target to improve the efficacy of the library comes at the expense of increasing library complexity, which in turn increases labor requirements as well as costs for reagents and sequencing. Because inactive constructs cannot be physically removed from the library, they serve as confounding factors during hit identification. While this can be partially mitigated with hits identification algorithms such as MAGecK^6^ and BAGEL^7^, there remains a need to further enhance CRISPR technology to improve library penetrance with smaller, more versatile pooled libraries. Several algorithms have been developed to aid in the design of sgRNAs with high efficiency guide ^4,8,9^ to minimize the number of constructs needed to yield robust results, but there remains a need for additional advancements to vastly decrease library size.

Similar to CRISPR/Cas9, CRISPR/Cpf1 (CRISPR from Prevotella and Francisella 1, or Cas12a) is a type II CRISPR system identified in the prokaryotic adaptive immune system that cleaves DNA target by small RNA guides. ^10^ It has been demonstrated that the Cpf1 orthologues LbCpf1 and AsCpf1 are highly specific, even more than SpCas9. ^11,12^ Moreover, Cpf1 is self-sufficient for multiplexed gene editing, unlike Cas9, which requires other Cas proteins and RNase III to process multiple guides in its native host. ^13^ Even though modified Cas9 multiplexing systems have been invented such as tRNA^14^, Cas6/Csy4^15^ and ribozyme^16^ aided multiplexing. Based on these properties, we reasoned a CRISPR/Cpf1 system may enable multiplexing of multiple guides targeting the same gene into a single lentiviral vector to generate high-efficiency pooled sgRNA libraries with substantially decreased complexity, therefore eliminate the trade-off paradox between library efficacy and library complexity.

To assess the performance of Cpf1 multiplexing, we generated our multiplexed AsCpf1 library targeting 342 “core-essential” genes and 345 “non-essential” genes, with three guides per gene ^3,4^(2061 guides, 687 constructs). Fitness change of knockouts of these genes are highly consistent across multiple cell lines, therefore making them “gold-standard” controls. To compare the screen performance of the multiplexed AsCpf1 library and conventional mono-cistronic CRISPR knock out libraries, we generated another two benchmark CRISPR libraries targeting the same group of genes: SpCas9-based mono-cistronic library (2061 guides, 2061 constructs) and AsCpf1-based mono-cistronic library (2061 guides, 2061 constructs). The design rules for SpCas9 and AsCpf1 guides are highly similar, despite the nuclease-specific requirements, such as different protospacer adjacent motifs (PAMs). The AsCpf1-based mono-cistronic and multiplexed libraries share identical guide sequences; however, the multiplexed AsCpf1 library has only a single construct harboring all three guides (***Fig 1a***).

**Fig. 1a.**
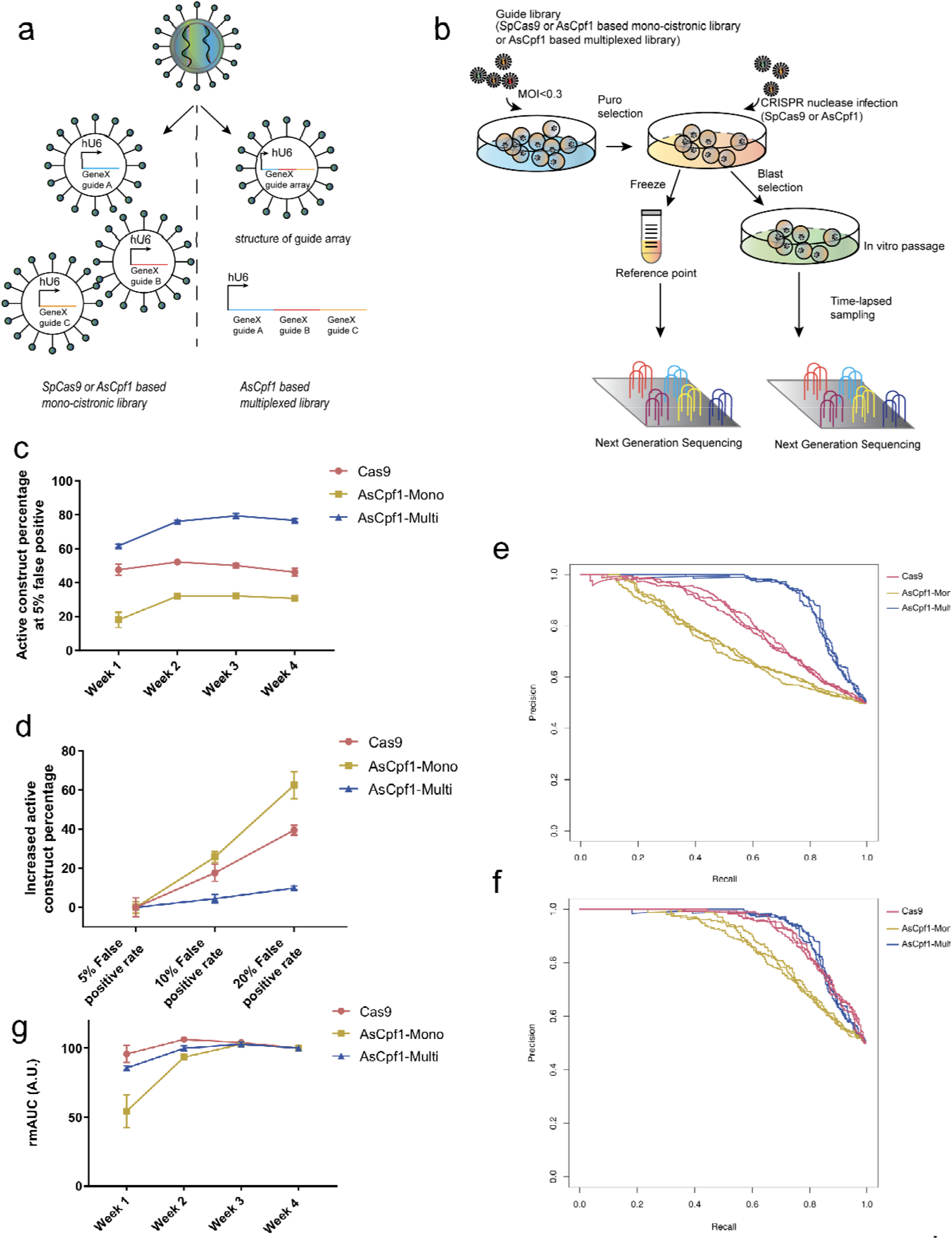
SpCas9 and AsCpf1 based guide libraries. In SpCas9 or AsCpf1 based mono-cistronic libraries, the basic unit is an SpCas9 or AsCpf1 guide, targeting one gene. Each gene has multiple guides dispersed in different lentivirus. In AsCpf1 based multiplexed library, the basic unit is an AsCpf1 guide array driven by one polymerase III promoter (e.g. human U6), targeting one gene. Each gene only has one corresponding guide array construct. However, each array contains multiple different guides targeting one gene. **Fig. 1b. Pooled library screen pipeline schematics.** K-562 were firstly infected with guide libraries (i.e. SpCas9 based mon-cistronic, AsCpf1 based mono-cistronic and AsCpf1 based multiplexed library) separately at low multiplicity of infection (<0.3) to ensure most infected cells were infected by one lentivirus. Two days post-infection, cells were selected with puromycin. After the non-infected cells were wiped out, a reference sample for all three libraries was collected and frozen for the following next-generation sequencing. The remaining cells were split into triplicates for each library and infected with lentivirus containing their corresponding CRISPR nuclease (i.e. for SpCas9 based library, SpCas9 nuclease was used. For AsCpf1 based libraries, AsCpf1 was used.). Cells were then selected with blasticidin to ensure SpCas9 and AsCpf1 were integrated and then screened in blasticidin and puromycin containing culture medium for 4 weeks. Samples were taken from each replicate every week to capture the screen dynamics. **Fig. 1c. The percentage of active construct for different libraries at the 5% false positive rate across 4 time points**. Data is based on log2 transformed fold change. Pink: Cas9 based mono-cistronic CRISPR library. Yellow: AsCpf1 based mono-cistronic CRISPR library. Purple: AsCpf1 based multiplexed CRISPR library. **Fig. 1d. The increased percentage of active construct for different libraries at the screen endpoint with a different controlled false positive rate.** Data is based on log2 transformed fold change. Pink: Cas9 based mono-cistronic CRISPR library. Yellow: AsCpf1 based mono-cistronic CRISPR library. Purple: AsCpf1 based multiplexed CRISPR library **Fig. 1e. The construct-level precision-recall curves for CRISPR libraries.** The construct-level precision-recall curves are based on the Bayes factor ranks for each guide at the endpoint. Each curve represents one biological replicate. Pink: Cas9 based mono-cistronic CRISPR library. Purple: AsCpf1 based multiplexed CRISPR library. Yellow: AsCpf1 based mono-cistronic CRISPR library. **Fig. 1f. The gene-level precision-recall curves for CRISPR libraries.** The gene-level precision-recall curves are based on the sum of Bayes factor ranks for all guides of the same gene at the endpoint. Each curve represents one biological replicate. Pink: Cas9 based mono-cistronic CRISPR library. Purple: AsCpf1 based multiplexed CRISPR library. Yellow: AsCpf1 based mono-cistronic CRISPR library. **Fig. 1g. The rmAUC curves of CRISPR libraries describing population dynamics.** rmAUC (ratio of the modified area under the curve) is calculated by (AUC_x_-0.498)/(AUC_end_-0.498) × 100%. AUC_x_: the area under the curve of construct-wise precision-recall curves for time point X. AUC_end_: the area under the curve of construct-wise precision-recall curves of endpoint. Pink: Cas9 based mono-cistronic CRISPR library. Yellow: AsCpf1 based mono-cistronic CRISPR library. Purple: AsCpf1 based multiplexed CRISPR library.

Benchmark screens employed K-562 cells separately infected with the pooled CRISPR libraries separately at a low multiplicity of infection (MOI). After puromycin selection, the corresponding CRISPR nuclease (AsCpf1 or SpCas9) was delivered by lentivirus transduction and blasticidin selection. Triplicate screens were conducted with cells grown for 4 weeks, and each replicate was sampled at intermediate time points to capture the dynamics of guide populations.

Screening performance was measured by separation of “core-essential” vs “non-essential” genes (***Fig 1b***). Interestingly, the common AsCpf1 variant used (human codon-optimized AsCpf1 with C-terminal nucleoplasmin bi-partite nucleus localization signal (NLS), herein AsCpf1-Nuc) ^11,17^ failed to show satisfactory gene-editing activity in the screen (***Supplementary Fig1a***), despite showing activity when tested with individual guides. (***Supplementary Fig1b***) However, an AsCpf1 variant bearing N-terminal 3x MYC-NLS and an optimized Kozak sequence (herein AsCpf1-3xMYC), effectively discriminated positive and negative control genes (***Supplementary Fig 1a, 1c***). When infected side-by-side with identical MOI, AsCpf1-3xMYC showed stronger expression and nuclear localization compared to AsCpf1-Nuc, suggesting that the amount of AsCpf1 in the nuclear fraction is critical for optimal gene editing efficiency. (**Supplementary *Fig 1d***)

All biological replicates for each of the three library screens correlated well, indicating good reproducibility (***Supplementary 2a, 2b, 2c***). An essential gene-targeting construct is considered active if it is more depleted compared to the non-essential gene targeting construct since it should have an anti-proliferative effect if it is active. To determine the percentages of active constructs among the three different libraries, we chose a false positive rate of 5% based on the log2 transformed fold change for each time point. The active construct percentage curve of the SpCas9-based mono-cistronic library was relatively flat across all four time points, with a mean value of 49.0% ± 2.9% active constructs. For AsCpf1-based libraries, the active construct percentage curve plateaued 2 weeks after the screen initiated with mean values of 31.7% ± 0.9% and 77.4% ± 1.4% active constructs for the mono-cistronic and multiplexed libraries, respectively (***Fig 1c***). The different shapes of the active construct percentage curves for SpCas9-and AsCpf1-based screens indicate different population temporal dynamics and knockout efficiency for the different CRISPR nucleases. Our data strongly suggest that SpCas9 is more active in mammalian cell gene knockout experiments compared with AsCpf1. However, multiplexing different guides targeting the same gene significantly increased the likelihood of gene knockout with AsCpf1. At endpoint, the percentage of active constructs in the AsCpf1 multiplexed library was only slightly increased compared to other libraries when we loose the FPR stringency from 5% to 20% (increased by 4.4% ± 1.9% at 10% FPR and 9.9% ± 0.8% at 20%FPR) (***Fig 1d***), indicating relatively low noise in the AsCpf1 multiplexed library screens.

To call out significantly depleted genes, we used an adapted Bayesian Analysis of Gene Essentiality (BAGEL) algorithm to analyze construct-level data. Based on the fold change in sgRNA abundance after knockout of each gene in the essential and non-essential training sets BAGEL uses a Bayesian model selection approach to classify a Bayes Factor (BF), which is the log2 likelihood of each gene belonging to either the essential gene distribution or non-essential gene distribution. Because BAGEL is designed for whole-genome CRISPR screens, we designed and utilized a version of BAGEL optimized for small library screens, “Low Fat BAGEL”. Low Fat Bagel generates BFs on a construct-level basis that is summed across guides to obtain a gene-level BF. For the AsCpf1-based multiplexed library, each gene has only one corresponding construct; therefore, its construct-level BF corresponds directly to its gene-level BF. To benchmark screen performance across the three libraries, precision-recall curves were plotted based on BFs. The precision-recall curves clearly showed that the SpCas9-based mono-cistronic screen (construct-wise area under the curve (AUC) 0.78 ± 0.01, gene-wise AUC 0.89 ± 0.01) outperformed the AsCpf1-based mono-cistronic screen (construct-wise AUC 0.70 ± 0.01, gene-wise AUC 0.82 ± 0.01) at both the construct (***Fig 1e***) and the gene level (***Fig 1f***). However, the AsCpf1-based multiplexed screen (construct-wise and gene-wise AUC 0.89 ± 0.00) performed similarly to the SpCas9 monocistronic library at the gene level, and it yielded a much stronger performance at the construct level, primarily due to lower active construct percentage in the SpCas9 screen, indicating a synergistic effect when guides were multiplexed together for the screening application (***Fig 1e,1f***). This is consistent with the reported synergistic effects observed in dCpf1-based activator-induced gene expression experiments ^18^ as well as with the increase in active constructs we observed in the context of multiplexed versus mono-cistronic AsCpf1 libraries.

To compare the rate of separation between essential and non-essential genes among the three library screens, we calculated the ratio of the modified area under the precision-recall curve (mAUC) of any given time point divided by the mAUC at endpoint (ratio of mAUC, rmAUC) (***Fig 1g***). As the area under the precision-recall curve (AUC) for this library would be 0.498 when there is no separation between essential and non-essential genes, the mAUC for any given time point was set to be its AUC minus 0.498. In accordance with ***Fig 1c***, the results suggest that separation between essential and non-essential genes in the AsCpf1-based mono-cistronic screens was much slower compared to SpCas9-based mono-cistronic and AsCpf1-based multiplexed screens. This might be the result of the relatively slower cleavage rate of AsCpf1 compared with SpCas9^19^, as we also saw a slightly slower separation between essential genes and non-essential genes in the AsCpf1 based multiplexed screen compared with SpCas9.

CRISPR/SpCas9 guide design has been optimized using empirical data from hundreds of screens^4,6,8,9^, but previous AsCpf1 guide optimization algorithms are largely based on a small number of surrogate reporters experiments ^17^. It is known that lentivirus has a preference on integration site^20^ and the chromosomal environment is one factor that influence CRISPR nuclease activity^21^. Thus, the gene editing process on surrogate reporters might not fully represent the true biological effect on endogenous loci editing. Our screen provided the first large-scale action-in-situ dataset to enable prediction of AsCpf1 guide preference based on functional screen data of endogenous loci. We used fold change information of essential gene-targeting guides in the mono-cistronic AsCpf1 library to calculate sequence preference, as effective guides should drop out more efficiently than ineffective guides. Even though all of the 342 genes are essential genes, the severity of their knockout phenotypes may differ. To avoid any sequence biases introduced by gene-specific effects, within the three guides in each gene, we classified the most depleted guide as the “high-performing” guide, and the least depleted guide as the “low-performing” guide. Using a scoring scheme similar to that of Hart et al. ^4^, the frequency of each nucleotide at each position of the 20-mer protospacer was calculated individually for the high-performing and low-performing guides. At each position, the nucleotide frequency of low-performing guide is subtracted from the high-performing guide to produce a table with subtracted frequencies for each nucleotide. This process was repeated across 100 bootstraps, and an aggregate average score table for each protospacer position was obtained. In agreement with the previous report ^17^, thymine (T) was strongly disfavored in position 1 in the protospacer, while guanine (G) and cytosine (C) were favored. We also identified a general trend of G disfavor from position 16 to position 20 in the protospacers. In addition, we found T was favored in positions 3,16, and 18, while C was favored in position 7 but disfavored in position 3 (***Figure 2a***). The score table obtained served as a metric to predict guide activity in terms of fold change: the sequence score for each guide is a sum of the nucleotide score at each position. Therefore, a guide with a zero sequence score indicates no similarity between either effective nor non-effective guides. We validated our prediction algorithm on the “median-performing” guides not used to develop the scoring algorithm – that is, the guide for each gene that was neither the highest nor lowest performing guide. Each median-performing guide was assigned a sequence score, and was classified with a prediction of “high-performing” or “low-performing” based on a guide score > 0 or ≤ 0, respectively. We then evaluated if the sequence score and predicted performance classification of each guide were indicative of fold change, and found a significant correlation between the guide score and guide performance (Spearman’s rho = −0.40, 95% confidence interval: (−0.34, −0.45), p=4.5 × 10^−40^) (***Figure 2b)***. The predicted high-performing guides were significantly more effective than low-performing guides, with a mean log2 fold change of −3.19 compared to −1.16 (t = −10.13, p = 4.9 × 10^−23^) (***Figure 2c)***.

**Fig. 2a.**
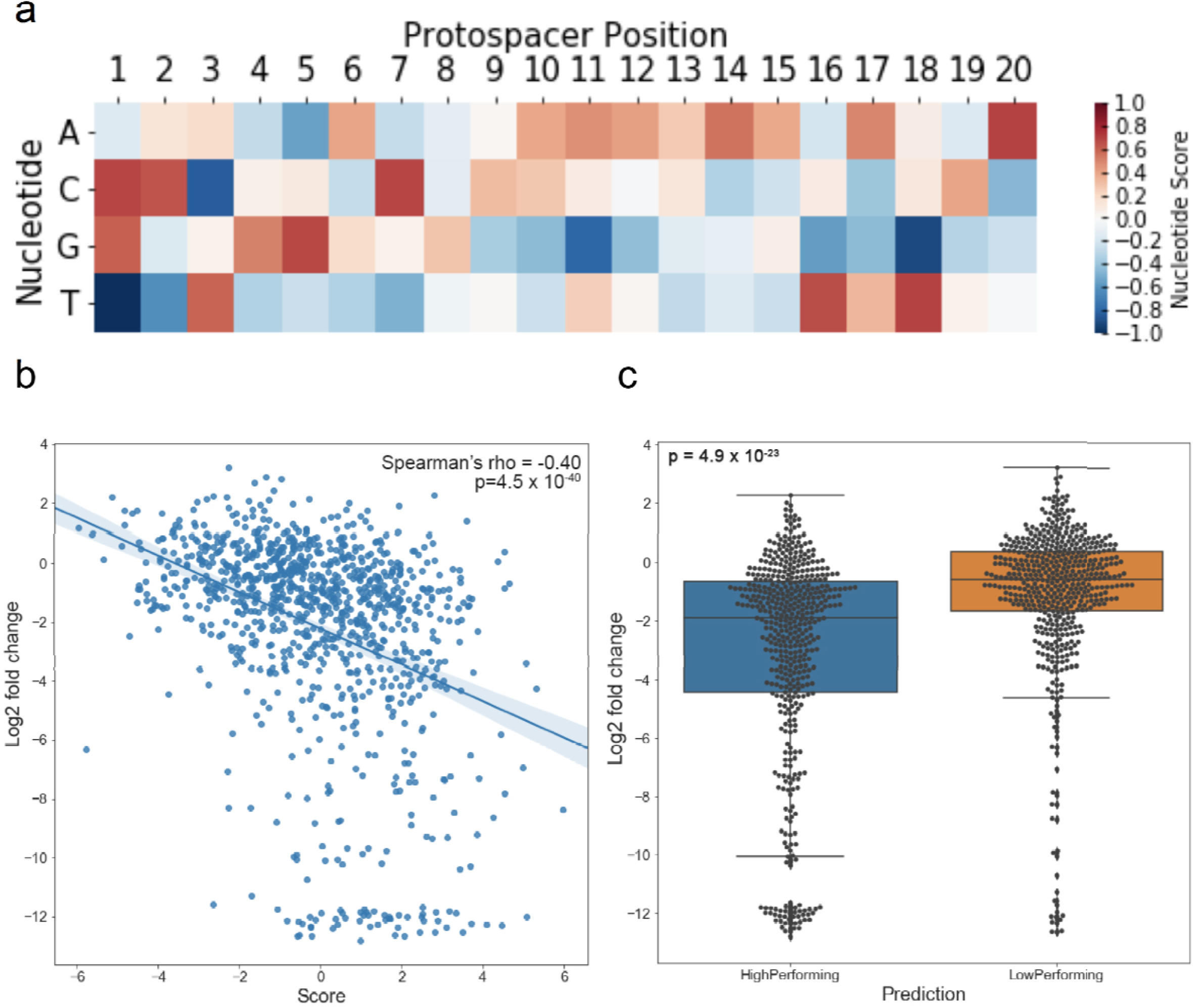
The scoring matrix for the prediction of AsCpf1 activity on endogenous loci. The screen data of mono-cistronic AsCpf1 library was used to generate the heat map of nucleotide preferences on protospacers. Positive value (in red tone) indicates a preference towards a nucleotide for a high performing guide while negative value (in blue tone) indicates a preference against a high performing guide. The position of the first nucleotide adjacent to PAM is define as position 1 for the protospacer. **Fig. 2b. Correlation between AsCpf1 guide prediction scores and Log2 transformed fold change** The predicted AsCpf1 guide scores for each validation guides were plotted against their log2 transformed fold change in the screen. (n=342) **Fig. 2c. Log2 transformed fold change of predicted high performing AsCpf1 guides and low performing guides** Guides were defined high performing if predicting score > 0 and low performing if predicting score ≤ 0. (n=342)

Based on our multiplexed AsCpf1 library strategy, we designed the smallest available CRISPR library targeting the entire human protein-coding genome, “Mini-human”. The guides for Mini-human were optimized based on activity scores derived from our screen dataset and further filtered for potential off-target effects. Because a previous analysis of published screens determined that four to six gRNAs per gene yields robust results when computational approaches to design sequence-optimized guides are employed ^4,6,9^, each construct in Mini-human contains up to 4 gene-targeting guides: 16393 gene-targeting constructs with 4 optimized guides, 584 gene-targeting constructs containing 3 optimized guides, and 55 non-targeting guide arrays as negative controls. This library is approximately one-fourth the size of the smallest currently available genome-wide CRISPR library and will be publicly available through Addgene.

In conclusion, with an AsCpf1 CRISPR-associated nuclease optimized for improved protein expression and nuclear localization, we have addressed the trade-off issue between library efficacy and library complexity for pooled libraries by multiplexing multiple guides in a single construct. We demonstrated that the multiplexed AsCpf1-3xMYC performed similarly well, if not better than, a conventional SpCas9-based mono-cistronic library, but with a significantly reduced library size. The new Mini-human library provides an invaluable tool for demanding functional genomics applications, especially *in vivo* pooled library screenings where library size is a concern.

## Acknowledgements

We thank Draetta lab members for helpful discussions. The human codon-optimized AsCpf1 was cloned from the kind gift plasmid SQT1659 from Keith Joung. Lenti-AsCpf1-Blast was a gift from Hyongbum Kim. LentiUniversal-Puro cannot be constructed without the generous gift LentiGuide-Puro from Feng Zhang. We would like to express our thanks to Florian Heigwer for the help of guide designing. T.H is supported by MD Anderson Cancer Center Support Grant P30 CA016672 (the Bioinformatics Shared Resource) and the Cancer Prevention Research Institute of Texas (CPRIT) grant RR160032.

## METHODS

### Cell Culture

K-562 cells (ATCC) were cultured in RPMI-1640(Hyclone) supplemented with 10% fetal bovine serum (Gibco, heat-inactivated), penicillin (100U/mL final concentration) and streptomycin (100U/mL). Lenti-X 293T(Clonetech) were cultured in DMEM (Hyclone) supplemented with 10% fetal bovine serum (Gibco, heat-inactivated), penicillin (100U/mL final concentration) and streptomycin (100U/mL).

### Plasmids

Human codon optimized AsCpf1 were PCR amplified with primers containing NLS signals in NEB Q5 hot start master mix from plasmid SQT1659 (Addgene# 78743). lenti-AsCpf1-Blast (Addgene# 84750) were digested with AgeI-HF(NEB) and BamHI-HF(NEB). NLS modified AsCpf1 and digested vector were ligated using Gibson Assembly.

### Immunoblotting

AsCpf1 antibody (Genetex, GTX133298, 1:250), Histone H3 antibody (CST, 9715,1:3000), Beta-tubulin (Sigma, T4026,1:3000) were incubated overnight at 4 °C. Anti-rabbit (CST,7074,1:2000) and anti-mouse secondary antibody (CST,7076,1:2000) were incubated for 3 hours at room temperature.

### Guide design

A total number of 342 core-essential genes and 345 non-essential genes were used to generate both Cas9 and AsCpf1 based libraries, with 3 guides per gene. Design of guides were accomplished by program CLD^22^. On-target selection rules were identical for both Cas9 and AsCpf1 except for the different requirements of PAM. Briefly, guides targeting the most transcripts and are closest to the first exon were prioritized. “NGG” PAM was used for Cas9 and “TTTV” was used for AsCpf1. Bowtie was used for “off-target” prediction. Mismatch tolerance of Cas9 was set to be up to 2 mismatches across 20bp spacer and up to 1 mismatch across PAM adjacent 18bp for AsCpf1. Any guides having over 2 predicted off-target sites across hg19 or targeting regions documented in dbSNP were excluded.1911 out of 2061 guides were predicted to be off-target free in Cas9 library and 1899 out of 2061 guides were in AsCpf1 library.

### Genome-wide “mini-human” multiplexed library design

Guides were identified by adapting the Cas9 library design algorithm developed by Hart et.al ^4^. Candidate guide sequences were obtained from exonic regions using hg38 and are filtered for homopolymers, and BsmBI restriction sites. Using Bowtie, we aligned the filtered candidate guides across the genome, allowing for one mismatch outside the “TTTV” PAM sequence. Guides with off-target matches in intronic or exonic regions were excluded, and the remaining guides were ranked based on the number of off-target matches in intergenic regions. A sequence score was assigned to each guide based on the score table presented in Figure 2a.

### Library screen

K-562 cell line was infected with each library at <0.3 multiplicity of infection at aimed coverage of at least 1000-fold. 48 hours post infection, normal cell culture medium was changed into cell culture medium containing 2 μg/ml puromycin. 4 days post infection, cells were infected with lentivirus generated from lentiCas9-Blast (Addgene #52962) and lentiAsCpf1-3xMYC correspondingly, ~1000X infected cells were isolated for each library for reference purpose. 6 days from the first infection, cells were further selected with 10 μg/ml blasticidin and split into triplicate for each group. Regular sampling was taken during the process of screening at each time point, cells were pelleted and frozen at −80 °C before further processing. At each sampling and screen process, cells were maintained at a minimum of 1000-fold complexity of the libraries per replicate.

### T7E1 assay

72 hours post transfection of guide and nuclease, cell genomic DNA was extracted with Qiagen DNeasy blood and tissue kit per manufacture’s protocol. Genomic DNA was used for the template of PCR with NEB Onetaq mastermix with standard buffer per manufacture’s protocol. The product was first denatured at 98°C for 5 minutes, then slowly annealed to 75°C at 1°C /s and eventually to 25°C at 0.1°C /s. 5U T7 Endonuclease I was used for digestion of less than 300 ng annealed product for 30 minutes. The digested product was subjected to 2% TAE agarose gel electrophoresis.

### Next generation sequencing

Genomic DNA were extracted from cell pellets using DNAzol (MRC Inc.) per manufacture’s protocol. Multiplexing barcodes and illumina sequencing adaptors were incorporated in the 1-step PCR with NEBNext Q5 Hot Start HiFi PCR Master Mix with following conditions: initial denaturing at 98°C for 1 min, denaturing at 98°C for 10s, annealing at 64°C for 20s, elongation at 72°C for 30s, final elongation for 2 min. PCR cycles for each sample were controlled to the minimal level where the target bands could be seen in 2% agarose TAE gel to ensure unbiased PCR amplification. Target bands were excised from the gel and purified with Freeze ‘N Squeeze™ DNA Gel Extraction Spin Columns (Bio-rad), quantified and pooled together. The pooled illumina library was then subjected to Nextseq550 high output sequencing. Reads were mapped with Bowtie and the mapped reads counts were used for further bioinformatics analysis.

### Low Fat BAGEL

The BAGEL computational framework estimates the distributions for the core essential and non-essential genes by bootstrapping the reference genes across 1000 permutations – where roughly 60% of the genes are randomly selected as a “training set”, and Bayes Factors (BF) for the remaining genes belonging to the “testing set” are calculated. The final BF for each gene is the average of the bayes factors obtained in the 1000 permutations. This method, while extremely robust in whole genome screens, results in overfitting of small library screens such as the benchmark screens we conducted as we are limited to 342 essential and 345 non-essential genes. To address this, we utilized a modified version of BAGEL suited for small libraries, termed Low Fat BAGEL. Rather than focusing on gene level BFs, we leveraged all available data by treating each sgRNA, for each gene, as an independent data point. Similar to BAGEL, 100 permutations are performed on the individual guides of the reference set, rather than on the level of each gene. The 500 guides are bootstrapped across 100 permutations, and the resulting BF for each guide computed is an average of all permutations. An aggregate BF for each gene is then obtained by summing the BFs of individual guides.

### Calculation of AUC for null distributions in the screen

When a screen with our benchmark libraries cannot distinguish essential gene targeting guides versus non-essential gene guides completely, the frequency distribution curves of these two categories overlap completely. At any given recall rate, the precision rate (true positive/all positive) should be a constant (i.e. the likelihood of an unknown guide X being an essential gene targeting guide, which is 342/ (342+345) =0.498). Therefore, for a null distribution, the AUC is 0.498. Lopes et.al for the detailed mathematical poof. ^23^

---

**Supp.1a.**
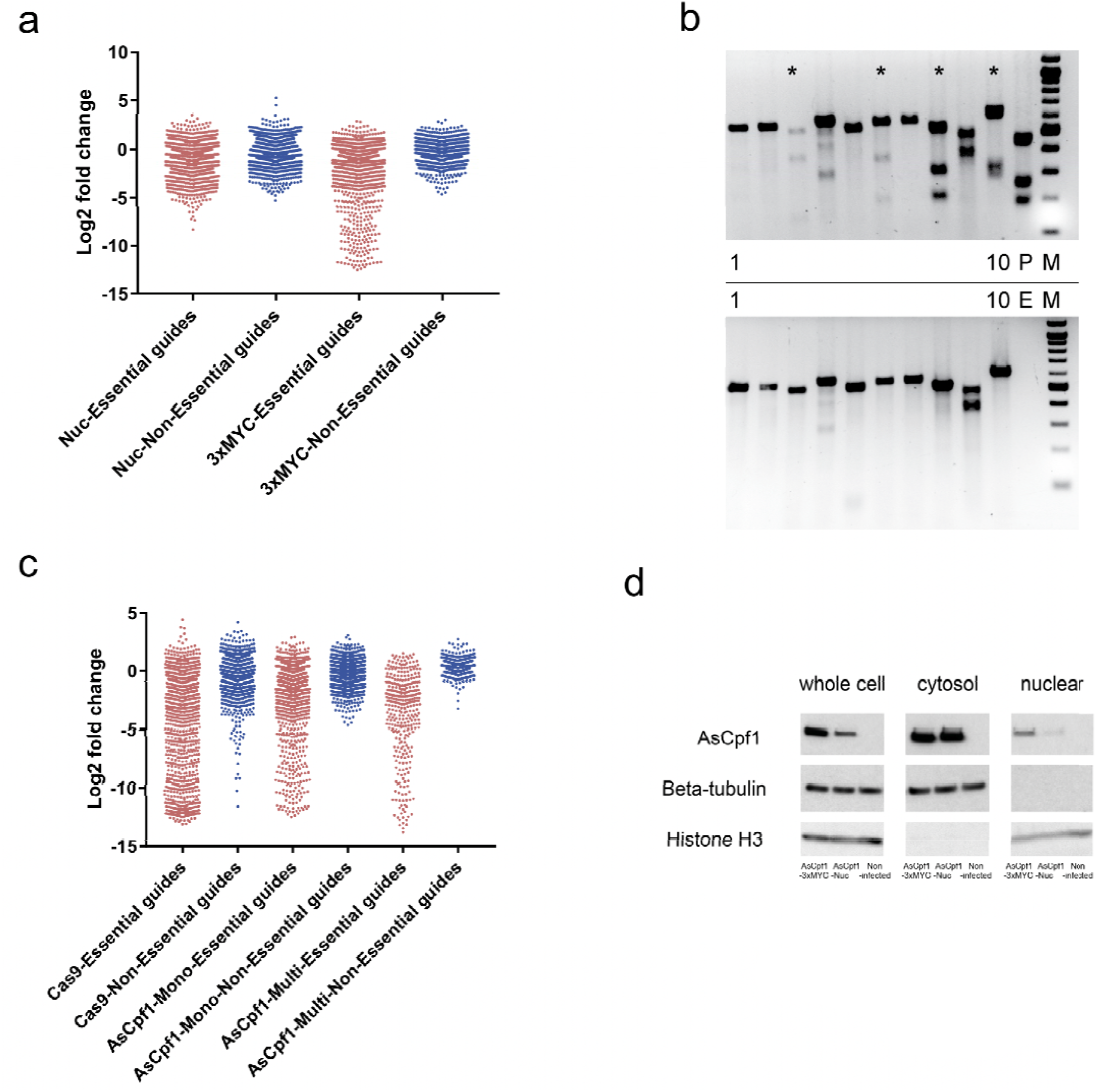
Log2 transformed guide fold change plot of AsCpf1-nuc variant and AsCpf1-3xMYC based screen. The AsCpf1-nuc variant based screen was conducted at the same controlled conditions with AsCpf1-3xMYC. Pink dot: the mean value of log2 transformed fold change of essential gene targeting guides at the endpoint. Purple dot: the mean value of log2 transformed fold change of non-essential gene targeting guides at the endpoint. **Supp**.**1b. T7E1 assay of 10 different loci transfected with AsCpf1-nuc and corresponding guides.** 293-T cells were transfected with AsCpf1-Nuc and guides in complexed with lipofectamine 3000. T7E1 assay was performed 72 hours post infection. Active guides are indicated by “*”. Upper panel: cells transfected with AsCpf1-nuc and guide. Lower panel: cells transfected with guides only. Lane1 to lane 10: T7E1 assay products targeting loci (from left to right) RESP18, NEUROD6, NAMPT, PRAMEF2, SLC22A9, COX7B2, TFAP2D, GUCY2F, ADAM30, and FNDC9. P: T7E1 assay product of DNMT1 edited allele as a positive control for T7 endonuclease activity. E: Empty lane. M: NEB 100bp DNA ladder. **Supp**.**1c. Log2 transformed guide fold change plot of AsCpf1-3xMYC and SpCas9 based screen.** Pink dot: the mean value of log2 transformed fold change of essential gene targeting guides at the endpoint. Purple dot: the mean value of log2 transformed fold change of non-essential gene targeting guides at the endpoint. **Supp**.**1d. Western blot of AsCpf1 variant.** K-562 cells were infected with AsCpf1-Nuc (the commonly used variant) and AsCpf1-3xMYC (Kozak sequence and nucleus localization signal optimized variant) virus at the same multiplicity of infection. After blasticidin selection, lysates of different cellular fraction were subjected to western blot. Beta-tubulin was the whole cell extract loading control and Histone H3 was the nucleus fraction lysate loading control. The AsCpf1-3xMYC variant showed stronger expression and nucleus localization.

**Supp.2a.b.c.**
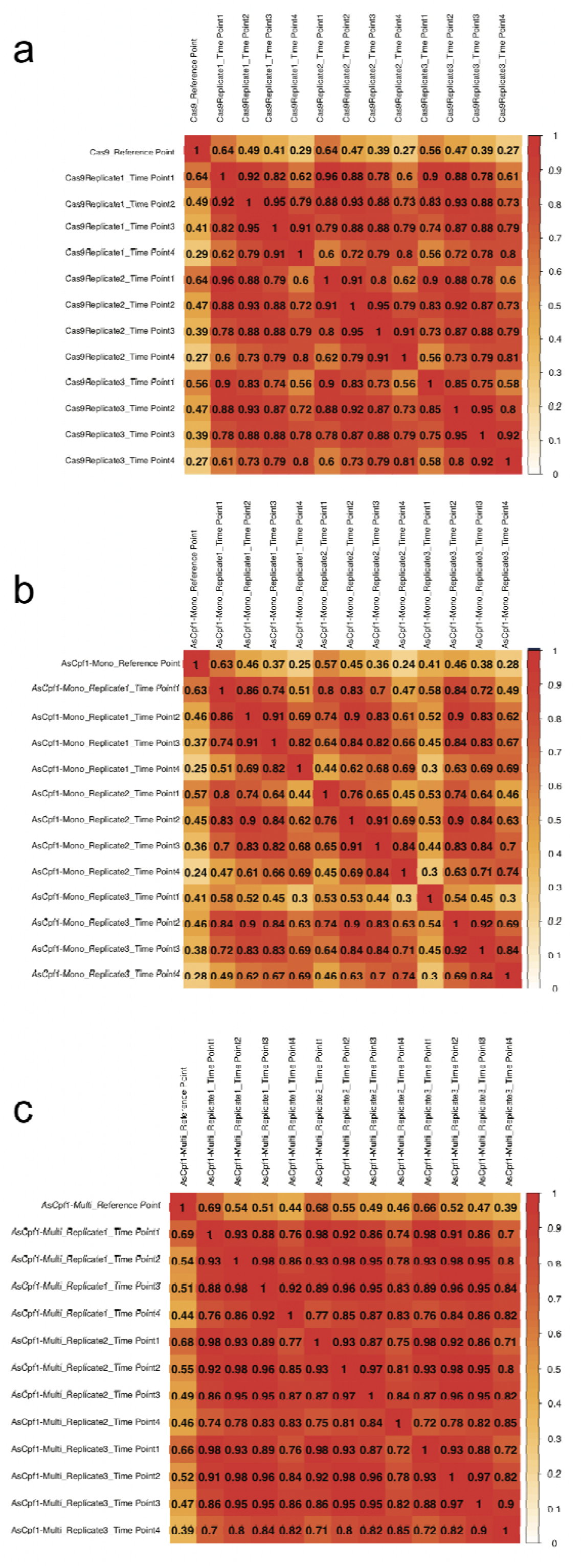
Pearson correlation coefficient matrices of all samples for SpCas9, AsCpf1 based library screen. Raw reads counts were used for the analysis.

**Supp.table.1** The protospacer sequences of randomly picked AsCpf1 guides and their corresponding genomic PCR primer sequences

**Supp.table.2** Library information for SpCas9 based benchmark library

**Supp.table.3** Library information for AsCpf1 based monocistronic benchmark library

**Supp.table.4** Library information for AsCpf1 based multiplexed benchmark library

**Supp.table.5** Raw reads counts for SpCas9 based benchmark library screen

**Supp.table.6** Raw reads counts for AsCpf1-3xMYC based monocistronic benchmark library screen

**Supp.table.7** Raw reads counts AsCpf1-3xMYC based multiplexed benchmark library screen

**Supp.table.8** Raw reads counts AsCpf1-Nuc based monocistronic benchmark library screen

**Supp.table.9** Library information for “Mini-Human” **Supp.text:** Sequence of AsCpf1-3xMYC vector

